# Two spatially distinct Kinesin-14 Pkl1 and Klp2 generate collaborative inward forces against Kinesin-5 Cut7 in *S. pombe*

**DOI:** 10.1101/205815

**Authors:** Masashi Yukawa, Yusuke Yamada, Tomoaki Yamauchi, Takashi Toda

**Affiliations:** Laboratory of Molecular and Chemical Cell Biology, Department of Molecular Biotechnology, Graduate School of Advanced Sciences of Matter, Hiroshima University, 1-3-1 Kagamiyama, Higashi-Hiroshima, Hiroshima 739-8530, Japan.

**Keywords:** fission, yeast, force, generation, kinesin, mitotic, bipolar, spindle, spindle, pole body

## Abstract

Kinesin motors play central roles in bipolar spindle assembly. In many eukaryotes, spindle pole separation is driven by Kinesin-5 that generates outward force. This outward force is balanced by antagonistic inward force elicited by Kinesin-14 and/or Dynein. In fission yeast, two Kinesin-14s, Pkl1 and Klp2, play an opposing role against Kinesin-5/Cut7. However, how these two Kinesin-14s coordinate individual activities remains elusive. Here we show that while deletion of either *pkl1* or *klp2* rescues temperature sensitive *cut7* mutants, only *pkl1* deletion can bypass the lethality caused by *cut7* deletion. Pkl1 is tethered to the spindle pole body, while Klp2 is localized along the spindle microtubule. Forced targeting of Klp2 to the spindle pole body, however, compensates for Pkl1 functions, indicating that cellular localizations, rather than individual motor specificities, differentiate between the two Kinesin-14s. Interestingly, human Kinesin-14/HSET can replace either Pkl1 or Klp2. Moreover, overproducing HSET induces monopolar spindles, reminiscent of the phenotype of Cut7 inactivation. Taken together, this study has uncovered the biological mechanism of how two different Kinesin-14s exert their antagonistic roles against Kinesin-5 in a spatially distinct manner.

**SUMMARY STATEMENT:** Proper force-balance generated by Kinesin-5 and Kinesin-14 is crucial for spindle bipolarity. Two fission yeast Kinesin-14s localize to different structures, thereby collaboratively producing inward forces against Kinesin-5-mediated outward force.

**Abbreviations used:**

GBP: GFP-binding protein
MWP complex: Msd1-Wdr8-Pkl1 complex
SPB: spindle pole body
ts: temperature sensitive
γ-TuC: the γ-tubulin complex

## INTRODUCTION

Identification of the canonical kinesin in squid axoplasm over 30 years ago (Vale et al., 1985) opened up the field towards our mechanistic understanding of microtubule-based, energy-dependent dynamic cellular processes. Subsequent molecular work has uncovered that kinesin proteins comprise a large family, which is categorized into three subfamilies, called N◻kinesin (N-terminal), M◻kinesin (middle) and C◻kinesin (C-terminal), depending on the position of microtubule-binding, ATPase domains within each molecule (Hirokawa et al., 2009; Lawrence et al., 2004). Phylogenetic analysis further divides these subfamilies into 14 groups (Kinesin-1 to -14); the majority of these groups belong to N-kinesins (Kinesins-1-12), M-kinesin includes Kinesin-13 and C-kinesin comprises Kinesin-14, which is further divided into two subgroups, Kinesin-14A and Kinesin-14B (Hirokawa et al., 2009). The most recent bioinformatic analysis indicates the existence of additional groups (Wickstead et al., 2010). In general, N-kinesins and C-kinesins possess microtubule plus end- and minus end-directed motilities respectively, while M-kinesins are microtubule depolymerases. As proposed originally (Vale et al., 1985), kinesin molecules act as cellular power plants by generating directional forces across microtubules through repeated ATPase cycles.

During mitosis, a number of distinct kinesin molecules engage with bipolar spindle assembly, thereby ensuring coordinated chromosome segregation; these kinesins are collectively called mitotic kinesins (Sharp et al., 2000b; Tanenbaum and Medema, 2010; Yount et al., 2015). In most, if not all, eukaryotes, a key mitotic kinesin for spindle formation is Kinesin-5 (budding yeast Cin8 and Kip1, fission yeast Cut7, *Aspergillus* BimC, *Drosophila* Klp61F, *Xenopus* Eg5 and human Kif11) (Blangy et al., 1995; Enos and Morris, 1990; Hagan and Yanagida, 1990; Heck et al., 1993; Le Guellec et al., 1991); it is arguably the only kinesin that is required for proper mitotic progression and cell viability. Kinesin-5 molecules form homo-tetramers, allowing them to crosslink and slide apart antiparallel microtubules, by which these motors generate outward pushing force onto spindle poles (Kapitein et al., 2005; Kashina et al., 1996). The inhibition of Kinesin-5 results in the appearance of monopolar spindles with duplicated, yet unseparated centrosomes/spindle pole bodies (SPBs) (Enos and Morris, 1990; Hagan and Yanagida, 1990; Heck et al., 1993; Hoyt et al., 1992; Mayer et al., 1999; Roof et al., 1992). The reasons for this monopolar spindle phenotype are deemed to be two-fold. One reason is rather passive; without outward pushing force produced by Kinesin-5, duplicated centrosomes/SPBs simply cannot split. The other reason is more active; in the absence of Kinesin-5, opposing motors such as Kinesin-14 or Dynein predominantly prevent centrosome/SPB separation by generating excessive inward force. As such, in many of the systems examined, inactivation of Kinesin-14 or Dynein rescues otherwise lethal Kinesin-5 inhibition (Bieling et al., 2010; Civelekoglu-Scholey et al., 2010; Ferenz et al., 2009; Gaglio et al., 1996; Gatlin et al., 2009; Mitchison et al., 2005; Mountain et al., 1999; O'Connell et al., 1993; Pidoux et al., 1996; Salemi et al., 2013; Saunders et al., 1997; Sharp et al., 2000a; Sharp et al., 1999; She and Yang, 2017; Tanenbaum et al., 2008; Tao et al., 2006; Walczak et al., 1998; Wang et al., 2015).

In many organisms, including fission yeast, human beings and plants, multiple Kinesin-14 proteins are encoded in the genome (fission yeast Pkl1 and Klp2, human HSET/KifC1, KifC2, KifC3 and Kif25 and *Arabidopsis* AKT and KCBP) (Decarreau et al., 2017; Noda et al., 2001; Pidoux et al., 1996; She and Yang, 2017; Troxell et al., 2001; Yamada et al., 2017). Even in budding yeast that contains the single Kinesin-14 Kar3, it forms two different complexes in a cell by interacting with distinct partners, Cik1 and Vik1 (Manning et al., 1999). Despite this multiplicity of Kinesin-14s, how individual molecules/complexes contribute to the generation of inward force against Kinesin-5-mediated outward force has yet to be satisfactorily addressed.

It would be worth pointing out that, although it is generally regarded that Kinesin-5 and Kinesin-14 are microtubule plus end- and minus end-directed motors respectively, a cohort of recent studies, mainly performed on fungal kinesins, have shown that some of these members undergo bi-directional movement in vitro and in some cases in vivo as well (Britto et al., 2016; Edamatsu, 2014; Gerson-Gurwitz et al., 2011; Popchock et al., 2017; Roostalu et al., 2011; Shapira et al., 2017).

In fission yeast, the lethality caused by temperature sensitive (ts) or even complete deletion of Kinesin-5-encoding *cut7* is suppressed by the additional deletion of Kinesin-14-encoding *pkl1* (Pidoux et al., 1996; Rodriguez et al., 2008; Syrovatkina and Tran, 2015; Troxell et al., 2001). We previously showed that Pkl1 forms a ternary complex with the other two proteins Msd1 and Wdr8 (Ikebe et al., 2011; Toya et al., 2007) (referred to as the MWP complex), which are transported toward the mitotic SPBs along spindle microtubules in a Pkl1-dependent manner. When Pkl1 reaches the SPB, it is tethered here in an Msd1-Wdr8-dependent manner as these two factors bind the SPB-localizing γ-tubulin complex (Vardy and Toda, 2000; Yukawa et al., 2015). SPB-tethered Pkl1 in turn generates inward force that antagonizes Cut7-mediated outward force.

In contrast to our understanding of Pkl1-mediated inward force aforementioned, the involvement of another Kinesin-14/Klp2 in force generation is less clearly defined. Although deletion of *klp2* reportedly rescues *cut7* ts mutants (Troxell et al., 2001), its underling mechanism remains undefined. It has been shown that Pkl1 and Klp2 perform different functions in both mitosis and meiosis (Grishchuk and McIntosh, 2006; Grishchuk et al., 2007; Troxell et al., 2001); however, the extent of functional similarities and diversifications between these two Kinesin-14s, in particular in terms of an antagonistic relationship with Kinesin-5/Cut7, has not been explored. Furthermore, how Pkl1 and Klp2 execute their individual roles in bipolar spindle formation in concert remains unclear.

In this study, we show that Pkl1 and Klp2 act to antagonize Kinesin-5/Cut7 in a functionally distinct manner. This difference is primarily attributable to their individual cellular localizations, rather than intrinsic motor specificities per se. Intriguingly, human Kinesin-14/HSET, when expressed in fission yeast, compensates for the loss of either Pkl1 or Klp2. We propose that collaborative actions of two spatially distinct Kinesin-14s are required for proper bipolar spindle assembly that antagonize Kinesin-5/Cut7.

## RESULTS

### Deletion of *klp2* rescues various *cut7* temperature sensitive mutants but not its complete deletion

Previous studies identified five independent *cut7* ts mutants (*cut7-21*, *-22*, *-23*, *-24* and *-446*) (Hagan and Yanagida, 1990, 1992; Pidoux et al., 1996), but no growth characteristics were compared, nor were mutated sites determined except for one allele (*cut7-22*, P1021S) (Olmsted et al., 2014). We first examined the growth properties of these ts mutants at various temperatures and found that individual mutants displayed different degrees of temperature sensitivity (Fig. S1A). Nucleotide sequencing analysis showed that each mutant contains a point mutation in different positions within Cut7; one in the N-terminal motor domain (*cut7-21*, M394T), another in the middle stalk region (*cut7-23*, S535P) and the remaining three near the C-terminal region (*cut7-22*, *-24* and *-446*, P1021S, V968A and I954T respectively) (Fig. S1B, note that P1021S in *cut7-22* resides within the conserved “BimC’ domain as previously pointed out) (Olmsted et al., 2014).

We then compared the ability of *pkl1*Δ, *klp2*Δ or *pkl1*Δ*klp2*Δ to suppress the temperature sensitivity of *cut7* alleles (*cut7-21*, *-22* and *-446*). As shown in Fig. 1A and B, either *pkl1*Δ or *klp2*Δ effectively, albeit not completely, rescued the ts phenotype of *cut7-21* and *cut7-22* at 36°C. Interestingly, growth of the most severe allele, *cut7-446*, was also ameliorated at 28°C (Fig. 1C). In general, we did not detect significant differences in suppression profiles between *pkl1*Δ and *klp2*Δ. The degree of suppression by double deletions differed depending on *cut7* ts allele used; a similar extent to each single deletion (*cut7-21*), a better or worse suppression (*cut7-22* or *cut7-446*, respectively). The reason for varied suppression profiles is currently unknown, though it could be related to the residual activity of each Cut7 ts mutant protein.

**Figure 1.**
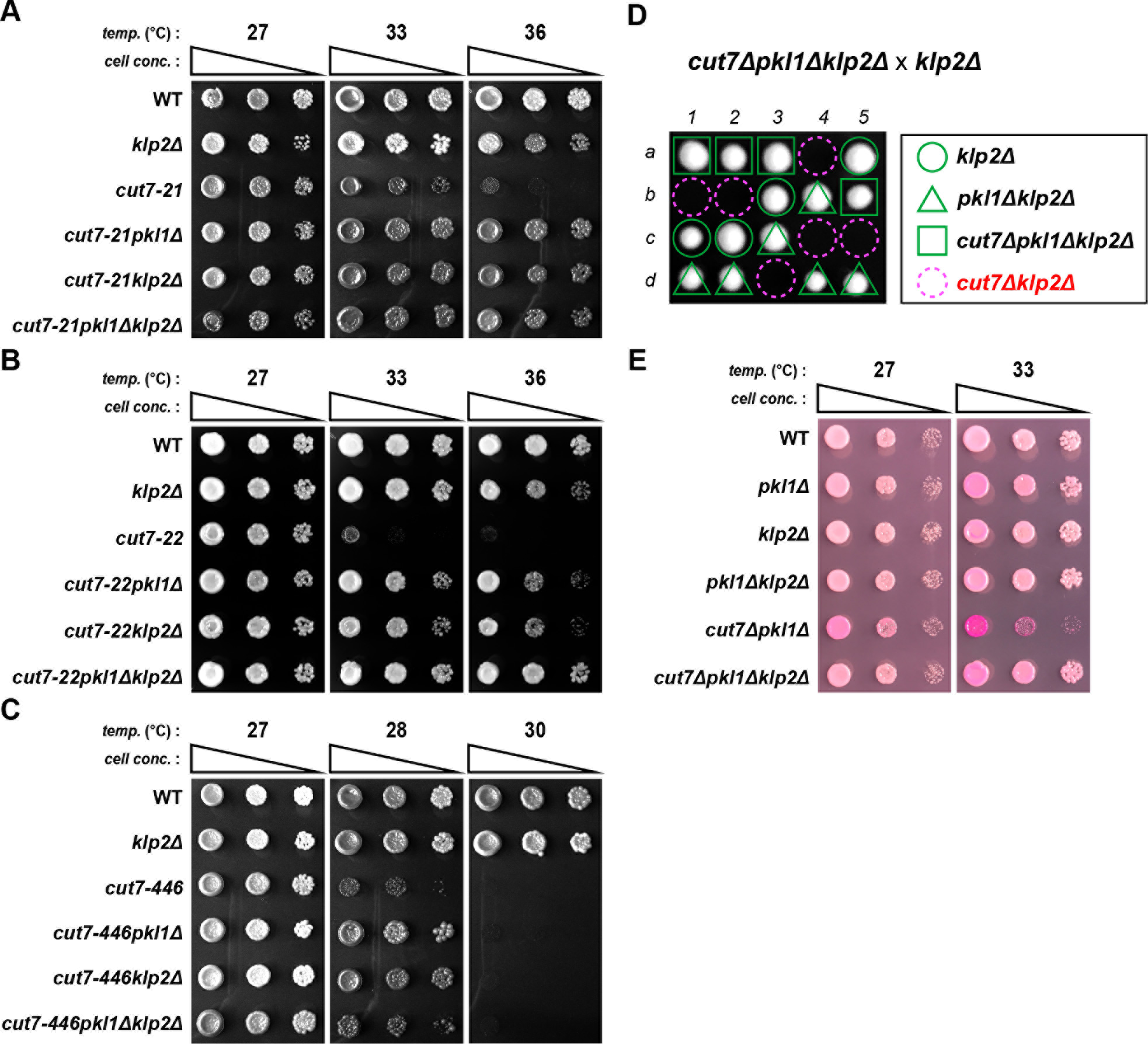
Absence of either Pkl1 or Klp2 is sufficient to rescue *cut7* ts mutants, but a single *klp2* deletion cannot compensate for the complete loss of Cut7. **(A-C)** Spot tests. Indicated strains were spotted onto rich YE5S agar plates and incubated at various temperatures for 3 d. 10-fold serial dilutions were performed in each spot. *cell conc*., cell concentration, *temp*., temperature. **(D)** Tetrad analysis. *cut7*Δ*pkl1*Δ*klp2*Δ and *klp2*Δ strains were crossed and allowed to mate and sporulate. Individual spores (*a-d*) in each ascus (*1-5*) were dissected on YE5S plates and incubated at 27°C for 3 d. Representative tetrad patterns are shown. Circles, triangles and squares with green lines indicate *klp2*Δ single mutants, *pkl1*Δ*klp2*Δ double mutants and *cut7*Δ*pkl1*Δ*klp2*Δ triple mutants, respectively. Assuming 2:2 segregation of individual markers allows the identification of lethal *cut7*Δ*klp2*Δ double mutants (indicated by dashed magenta circles). **(E)** Spot tests. Indicated strains were spotted onto rich YE5S containing Phloxine B (a red dye that stains dead or sick cells) (Moreno et al., 1991) at 27°C or 33°C for 3 d. *cell conc*., cell concentration, *temp*., temperature.

Next, we asked whether *klp2*Δ suppresses the lethality of *cut7*Δ. Previous work showed that *pkl1*Δ rescues *cut7*Δ (Olmsted et al., 2014; Syrovatkina and Tran, 2015). Tetrad dissection clearly showed that unlike *pkl1*Δ, *klp2*Δ was incapable of rescuing complete deletion of *cut7* (Fig. 1D). As *cut7*Δ*pkl1*Δ cells are viable, we created a *cut7*Δ*pkl1*Δ*klp2*Δ triple mutant strain and examined growth properties. As shown in Fig. 1E, *cut7*Δ*pkl1*Δ*klp2*Δ cells grew much better, particularly at 33°C, the temperature at which *cut7*Δ*pkl1*Δ double mutants exhibited retarded growth. Taken together, these results indicate that Klp2 antagonizes Cut7 additively with Pkl1; however, unlike Pkl1, *klp2* deletion on its own is not sufficient to bypass the requirement of Cut7. Hence, the functional contribution to the generation of inward force elicited by Pkl1 and Klp2 is not equal.

### *cut7-22klp2*Δ cells display two types of mitotic profiles

To investigate and visualize the impact of *klp2*Δ on mitotic progression of *cut7-22* ts cells, we observed the dynamic behavior of spindle microtubules and SPBs in wild-type, *cut7-22pkl1*Δ, *cut7-22klp2*Δ and *cut7-22pkl1*Δ*klp2*Δ cells that were incubated at 36°C (note that a single *cut7-22* strain cannot be used as cells are arrested in early mitosis with monopolar spindles) (Hagan and Yanagida, 1990). All strains contained mCherry-Atb2 (α2-tubulin, a microtubule marker) and GFP-Alp4 (a constitutive component of the γ-tubulin complex, an SPB marker) (Toda et al., 1984; Vardy and Toda, 2000). We found that, as previously reported (Yukawa et al., 2015), spindle length of *cut7-22pkl1*Δ cells at anaphase onset (defined as the timing when spindles start to elongate towards both cell ends after metaphase) (Nabeshima et al., 1998) was significantly shorter than that of wild type cells (0.96 ± 0.34 μm, n=32 vs 1.90 ± 0.32 μm, n=26 for wild type; Fig. 2A,B) and cells remained at a pre-anaphase stage for a much longer period of time (16 ± 7 min, n=32 vs 7 ± 2 min, n=26 for wild type; Fig. 2A,C). In addition, spindle elongation rate during anaphase B was considerably slower (0.53 ± 0.14 μm/min, n=32 vs 0.72 ± 0.10 μm/min, n=26 for wild type; Fig. 2A,D). Therefore, mitotic progression of *cut7-22pkl1*Δ cells is compromised.

**Figure 2.**
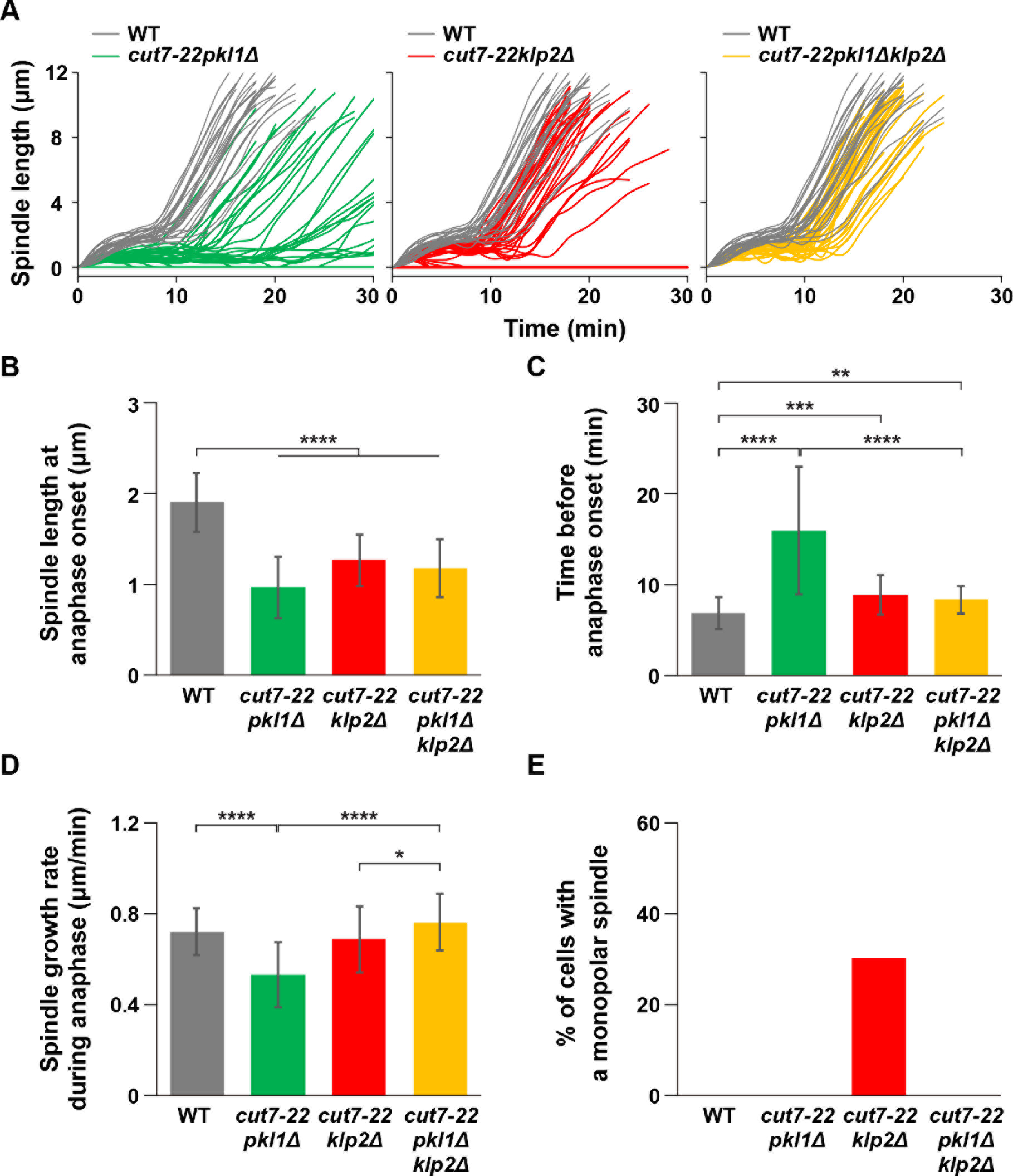
Mitotic profiles of *cut7-22klp2*Δ cells consist of two distinct populations. **(A)** Profiles of mitotic progression in *cut7-22pkl1*Δ (green line, n=32), *cut7-22klp2*Δ (red line, n=43) or *cut7-22pkl1*Δ*klp2*Δ cells (yellow line, n=32). Each strain contains a an SPB maker (GFP-Alp4) (Vardy and Toda, 2000) and a microtubule marker (mCherry-Atb2) (Toda et al., 1984). Cells were grown at 36°C for 2 h and live imaging performed thereafter. Changes of the inter-SPB distance were plotted against time. In each panel, patterns of wild-type cells are plotted for comparison (gray line, n=26). **(B)** Spindle length at anaphase onset. **(C)** The time between the initiation of SPB separation and onset of anaphase B. **(D)** Spindle growth rate during anaphase B. **(E)** The percentage of cells containing monopolar spindles (n>25). Data are given as means ± SD; *, P < 0.05; **, P < 0.01; ***, P < 0.001; ****, P < 0.0001 (two-tailed unpaired Student’s *t*-test).

The *cut7-22klp2*Δ double mutant cells displayed two distinct phenotypes with regards to mitotic parameters. In the majority of cells (~70%, 30 out of 43), mitotic progression was similar to that of *cut7-22pkl1*Δ cells, although less compromised. For instance, spindle length at anaphase onset was 1.26 ± 0.28 μm (n=30) (0.96 ± 0.34 min, n=32 and 1.90 ± 0.32 μm, n=26 for *cut7-22pkl1*Δ and wild type cells respectively, Fig. 2B) and the duration of the pre-anaphase period was 9 ± 2 min (n=30) (16 ± 7 min, n=32 and 7 ± 2 min, n=26 for *cut7-22pkl1*Δ and wild type cells respectively, Fig. 2C). The spindle elongation rate during anaphase B was 0.69 ± 0.15 μm/min (n=30), which is marginally slower than that of wild type cells (0.72 ± 0.10 μm/min, n=26 Fig. 2D). In sharp contrast, the remaining ~30% of *cut7-22klp2*Δ cells (13 out of 43) exhibited the persistent monopolar spindle phenotype for more than 30 min (Fig. 2E), which is reminiscent of *cut7-22* cells and is not observed in *cut7-22pkl1*Δ cells. Why these two distinct phenotypes emerge in *cut7-22klp2*Δ has not been explored further at the moment. It is possible that the hypomorphic *cut7-22* mutation does not lose gene function completely at the restrictive temperature; instead, this mutant protein may contribute to spindle formation to some extent, leading to the appearance of two differed phenotypic outcomes.

Intriguingly, mitotic profiles of the *cut7-22pkl1*Δ*klp2*Δ triple mutant cells were improved compared with those of either *cut7-22pkl1*Δ or *cut7-22klp2*Δ cells, although they were still not those of wild type cells. Most remarkably, cells displaying monopolar spindles (observed in 30% of *cut7-22klp2*Δ) were no longer present (Fig. 2A). Furthermore, the spindle elongation rate during anaphase B became 0.76 ± 0.13 μm/min (n=32) (Fig. 2D), which is faster than either *cut7-22pkl1* (0.53 ± 0.14 μm/min, n=32) or *cut7-22klp2*Δ cells (0.69 ± 0.15 μm/min, n=30). Spindle length at anaphase onset (1.17 ± 0.32 μm, n=32) and the duration of a pre-anaphase stage (8 ± 2 min, n=32) are still compromised compared to wild type cells (1.90 ± 0.32 μm and 7 ± 2 min, n=26, respectively) and similar to those of *cut7-22klp2*Δ (1.26 ± 0.28 μm and 9 ± 2 min, n=30 respectively). This result indicates that abnormal properties of mitotic progression of *cut7-22pkl1*Δ or *cut7-22klp2*Δ cells are partly ascribable to the presence of Klp2- or Pkl1 -mediated inward force respectively. This accounts for the foregoing results that growth of *cut7-22pkl1*Δ*klp2*Δ cells was noticeably improved compared with that of *cut7-22pkl1*Δ or *cut7-22klp2*Δ cells (see Fig. 1A).

### Triply deleted *cut7*Δ*pkl1*Δ*klp2*Δ cells display ameliorated, but sill compromised mitotic progression

We next examined mitotic profiles of *cut7*Δ*pkl1*Δ*klp2*Δ triple mutant cells and compared them with those of *cut7*Δ*pkl1*Δ cells (note that *cut7*Δ*klp2*Δ cells are inviable, see Fig. 1D). As shown in Fig. 3, we found that *cut7*Δ*pkl1*Δ*klp2*Δ cells display a substantial improvement in mitotic progression compared to that observed in *cut7*Δ*pkl1*Δ cells (Fig. 3A). For instance, spindle length at anaphase onset was 1.07 ± 0.29 μm (n=17) (1.01 ± 0.27 μm, n=15 for *cut7*Δ*pkl1*Δ, Fig. 3B) and the period of a pre-anaphase stage was 15 ± 3 min (n=17) (24 ± 4 min, n=15 for *cut7*Δ*pkl1*Δ, Fig. 3C). Furthermore, the rate of anaphase spindle elongation was also improved (0.36 ± 0.07 μm/min, n=15 vs 0.46 ± 0.08 μm/min, n=17 for *cut7*Δ*pkl1*Δ and *cut7*Δ*pkl1*Δ*klp2*Δ cells, respectively, Fig. 3D). These results indicate that Klp2 collaborates with Pkl1 in the generation of inward force that antagonizes outward force exerted by Cut7. It is of note that the mitotic profiles of these triply deleted cells are still not the same as those of wild-type cells (Fig. 3A), implying that pathways responsible for bipolar spindle assembly in the absence of Kinesin5- and Kinesin14 are less proficient in this role (Rincon et al., 2017; Yukawa et al., 2017).

**Figure 3.**
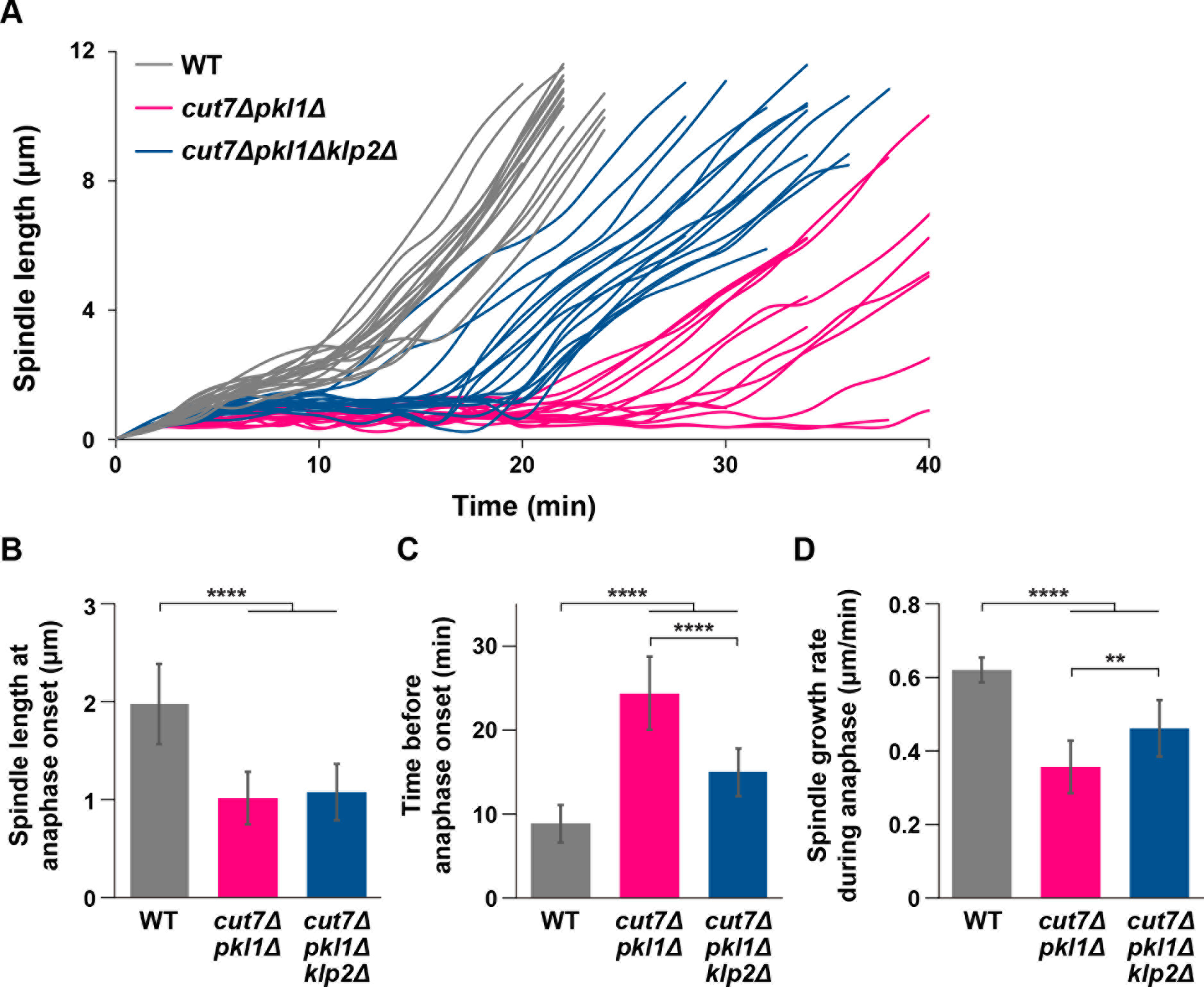
Deletion of *klp2* alleviates compromised mitotic progression and spindle abnormalities of *cut7*Δ*pkl1*Δ cells. **(A)** Profiles of mitotic progression in wild-type (gray line, n=17), *cut7*Δ*pkl1*Δ (red line, n=15) or *cut7*Δ*pkl1*Δ*klp2*Δ cells (blue line, n=17). Each strain contains fluorescent markers for the nuclear membrane (Cut11-GFP) (West et al., 1998), SPB (GFP-Alp4) and tubulin (mCherry-Atb2). Cells were grown at 30°C. Changes of the inter-SPB distance were plotted against time. **(B)** Spindle length at anaphase onset. **(C)** The time between the initiation of SPB separation and onset of anaphase B. **(D)** Spindle growth rate during anaphase B. Data are given as means ± SD; **, P < 0.01; ***, P < 0.001; ****, P < 0.0001 (two-tailed unpaired Student’s t-test).

### Klp2 is dispensable for anchoring the spindle microtubule to the mitotic SPB, but it secures spindle anchorage in collaboration with Pkl1

Pkl1 forms the ternary MWP complex containing Msd1 and Wdr8 and is thereby tethered to the mitotic SPB, which in turn anchors the minus end of the spindle microtubule to the SPB (Yukawa et al., 2015). This anchorage is required for generating inward force, as in the absence of Pkl1 (Msd1 or Wdr8 as well), free minus ends of the spindle microtubules physically push the nuclear membrane due to Cut7-mediated outward force, leading to the distortion of the nuclear envelope with spindle protrusions. We examined whether such similar protrusions were observed in the absence of Klp2. Despite careful inspection, we did not detect the characteristic protruding spindles or deformed nuclear membranes in *klp2*Δ cells (Fig. 4A).

**Figure 4.**
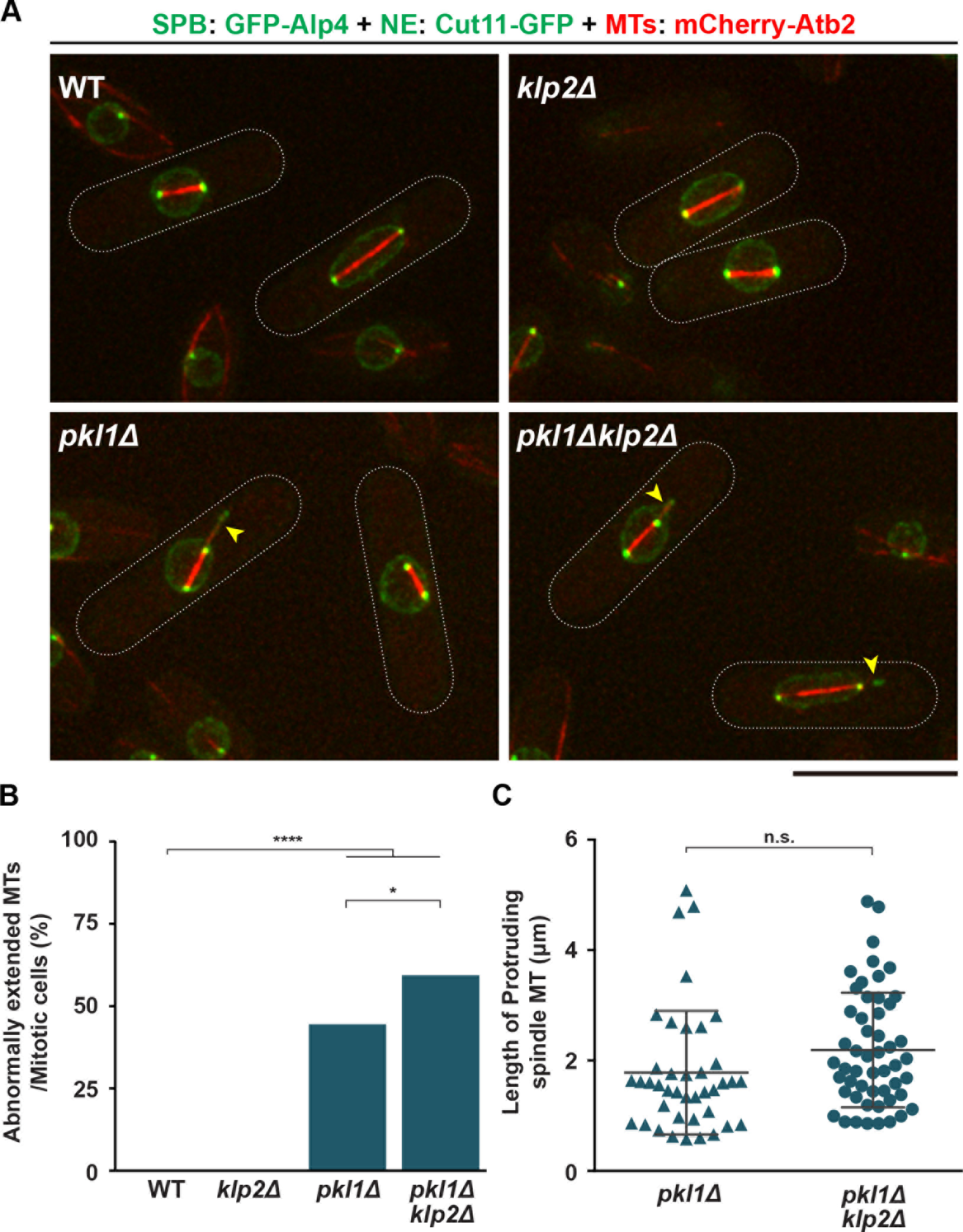
Klp2 is not required for but acts collaboratively with Pkl1 in anchoring the spindle microtubule to the mitotic SPB. **(A)** Representative images showing mitotic cells of indicated strains. Each strain contains fluorescent markers for the nuclear membrane (Cut11-GFP), SPB (GFP-Alp4) and tubulin (mCherry-Atb2). Deformed nuclear membranes encompassing protruding spindles are shown with arrowheads. Cells were grown at 27°C. Scale bar, 10 μm. **(B)** Quantification of spindle protrusions. In each strain, n>30 mitotic cells were examined. All p-values are derived from the two-tailed χ^2^ test (*, P < 0.05; ****, P < 0.0001). **(C)** Length distributions of protruding spindles. In either strain, n>40 mitotic cells with protruding spindles were observed and the length of protrusion was measured. Data are given as means ± SD; n.s., not significant (P=0.07, two-tailed unpaired Student’s *t*-test).

When combined with *pkl1*Δ, however, *pkl1*Δ*klp2*Δ cells exhibited more frequent spindle protrusions (Fig. 4A,B, n>30). The length of protrusions was also increased, though only marginally (Fig. 4C, n>40). This indicates that Klp2 plays an inhibitory role in spindle protrusion in collaboration with Pkl1. Hence, we conclude that provided Pkl1 is present, the minus end of the spindle microtubules is anchored to the SPB, which is Klp2-independent. We envisage that the residual inward force generated by Pkl1 at the SPB underlies the lethality of *cut7*Δ*klp2*Δ cells. This accounts for the emergence of monopolar spindles in some proportion of *cut7-22klp2*Δ cells (Fig. 2A,E) (see Discussion).

### Forced tethering of Klp2 to the SPB compensates for defective anchoring of the spindle microtubule in *pkl1*Δ cells

To explore the underlying mechanism of the functional differentiation between Pkl1 and Klp2, we first examined their cellular localization during mitotic progression. As previously shown (Yukawa et al., 2015), Pkl1 is predominantly localized to the SPB and only faintly on the spindle microtubule (Fig. 5A). In sharp contrast, Klp2 is localized mainly along the spindle microtubule often in a punctate manner; importantly, unlike Pkl1, Klp2 did not accumulate at the SPB (Fig. 5B). Some of the dots on the spindle microtubule may correspond to the kinetochores as previously noted (Troxell et al., 2001). Thus, mitotic localization patterns of these two Kinesin-14s are different.

**Figure 5.**
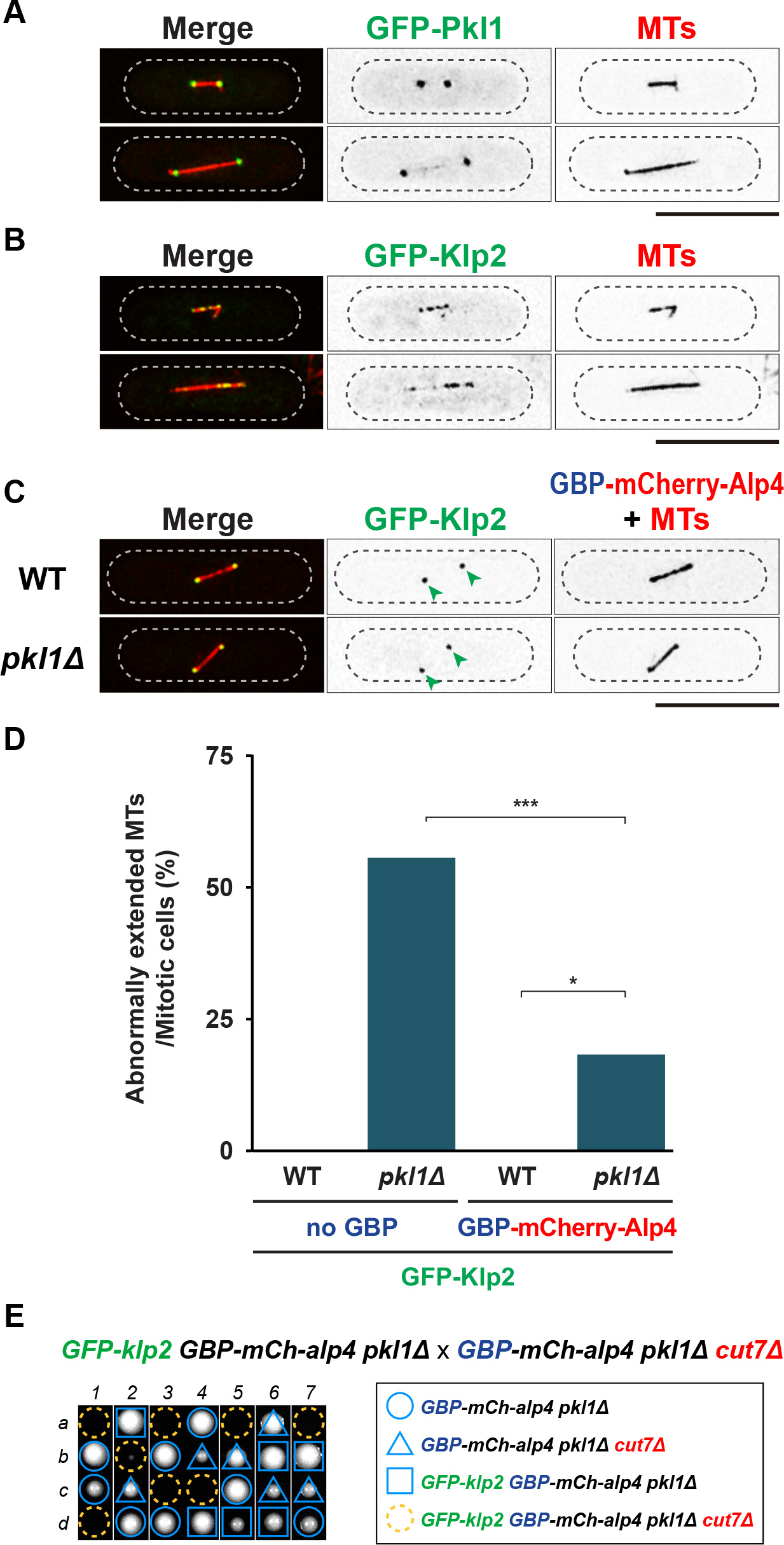
Distinct localization profiles of Pkl1 and Klp2 and a functional exchangeability by tethering of Klp2 to the SPB. **(A, B)** Representative mitotic images of GFP-Pkl1 or GFP-Klp2. Each strain contains a microtubule marker (mCherry-Atb2; red) and GFP-Pkl1 (**A**, green) or GFP-Klp2 (**B**, green). Cells were grown at 27°C. The cell peripheries are outlined with dotted lines. Scale bars, 10 μm (top). **(C)** Forced tethering of GFP-Klp2 to the SPB in wild-type (top row) or *pkl1*Δ cells (bottom row). GFP-Klp2 was recruited to the SPB using GBP-mCherry-Alp4, which is a constitutive component of the γ-tubulin complex (Vardy and Toda, 2000). Representative images of GFP-Klp2 during mitosis are shown. SPB-tethered GFP-Klp2 signals are marked with arrowheads. Scale bar, 10 μm. **(D)** Rescue of spindle anchoring defects in *pkl1*Δ cells by forced tethering of Klp2 to the SPB. >40 mitotic cells were observed in each strain. All p-values are derived from a two-tailed χ^2^ test (*, P < 0.05; ***, P < 0.001). **(E)** Tetrad analysis. A strain containing GFP-tagged Klp2 was crossed with *cut7*Δ and allowed to mate and sporulate (note that both strains are *GBP-mCherry-alp4pkl1*Δ). Individual spores (*a-d*) in each ascus (*l-7*) were dissected on YE5S plates and incubated at 27°C for 3 d. Representative tetrad patterns are shown. Circles, triangles and squares with blue lines indicate *GBP-mCherry-alp4pkl1*Δ, *GBP-mCherry-alp4cut7*Δ*pkl1*Δ and *GFP-klp2GBP-mCherry-alp4pkl1*Δ respectively Assuming 2:2 segregation of individual markers allows the identification of the remaining lethal segregants *GFP-klp2GBP-mCherry-alp4cut7*Δ*pkl1*Δ (indicated by dashed yellow circles).

We next sought to address whether the functional differences between Pkl1 and Klp2, in particular mechanisms for inward force generation, are derived from individual motor specificities or cellular location. For this purpose, we adopted the GFP entrapment strategy based upon the implementation of GFP-binding protein (GBP) (Rothbauer et al., 2008). We forced GFP-Klp2 to localize to the SPB using GBP-tagged Alp4, which is a constitutive component of the γ-tubulin complex (Vardy and Toda, 2000). Tethering of GFP-Klp2 to the SPB was successful and this localization was not dependent upon Pkl1 (Fig. 5C). We then assessed the emergence of the spindle protrusion phenotype in a *pkl1*Δ strain. Intriguingly, while *pkl1*Δ cells containing only GFP-Klp2 displayed a high frequency of spindle protrusions (~55%, n=65, Fig. 5D), in those containing both GFP-Klp2 and GBP-Alp4, the frequency was reduced to ~18% (n=44, Fig. 5D). Hence, targeting Klp2 to the SPB substantially mitigated anchoring defects resulting from *pkl1* deletion, albeit not completely.

We then asked whether SPB-tethered Klp2 is capable of genetically substituting for Pkl1 in the absence of Cut7. Tetrad analysis showed that *cut7*Δ*pkl1*Δ strains are inviable when they contain GFP-Klp2 and GBP-Alp4, indicating that SPB-localizing Klp2 plays an analogous role to Pkl1 (Fig. 5E). These results strongly suggest that it is the spatial regulation of the proteins rather than the intrinsic motor specificity that determines the functional differentiation between these two Kinesin-14s.

### Human Kinesin-14/HSET functionally compensates for the loss of either Pkl1 or Klp2

Human Kinesin-14/HSET opposes the activity of Eg5/Kif11 by producing inward force (Mountain et al., 1999). It has also been shown that HSET, when expressed in fission yeast, can rescue the loss of Pkl1 (Simeonov et al., 2009). However, whether HSET can functionally replace Klp2 is not known. Accordingly, we examined in more detail the functional exchangeability between HSET and fission yeast Kinesin-14s to explore the evolutionary conservation and diversification of Kinesin-14s.

Expressing the *HSET* gene tagged with *GFP* in fission yeast using the thiamine-repressible *nmt41* promoter (Basi et al., 1993) on plasmids showed that under repressed conditions, HSET was localized to the spindle microtubule, apparently uniformly. We did not detect the regional accumulation of GFP-HSET at or in proximity to the SPBs (Fig. 6A). We subsequently wondered whether HSET could generate inward force in place of Pkl1 or Klp2 by introducing it to *cut7-22pkl1*Δ or *cut7-22klp2*Δ respectively. If HSET were functionally exchangeable, these cells would display the ts phenotype. The results showed that HSET could compensate for the loss of either Pkl1 or Klp2 (Fig. 6B). We also asked whether HSET suppressed the protruding spindle phenotype of *pkl1*Δ cells. No significant improvement was detected; ~53% cells showed protruding spindles (n>25). Thus, although HSET can generate inward force that antagonizes Cut7-driven outward force in place of either Pkl1 or Klp2, HSET molecules failed to fulfill the spindle-anchoring role for Pkl 1; phenotypic suppression was only partial.

**Figure 6.**
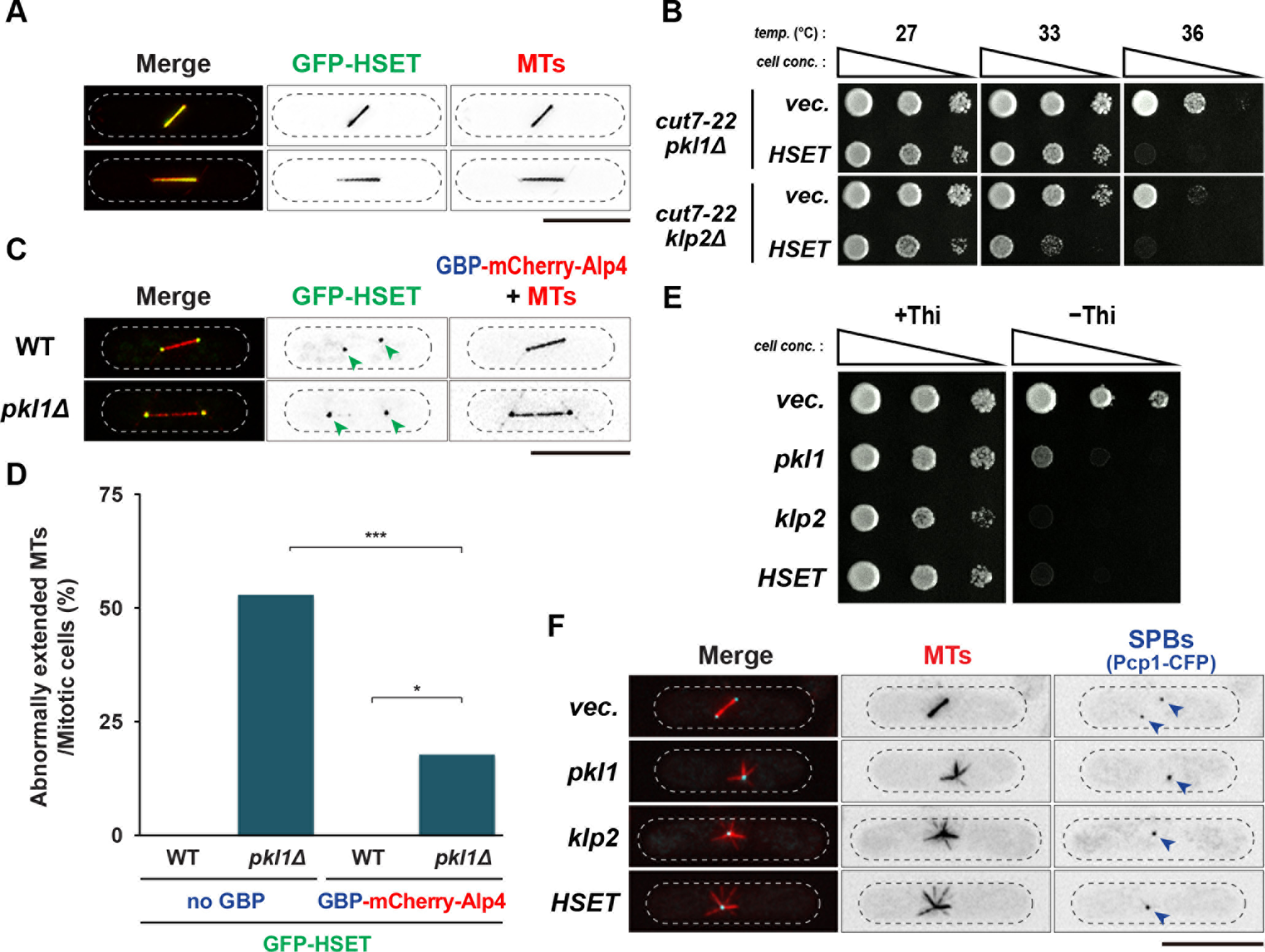
Functional suppression of loss of Pkl1 or Klp2 by human HSET. **(A)** Representative images of GFP-HSET during mitosis are shown. An episomal plasmid pREP41-GFP-HSET (green) was introduced into cells that contain a microtubule marker (mCherry-Atb2; red). Cells were grown in liquid minimal medium containing thiamine (repressed condition) at 30°C. Scale bar, 10 μm. **(B)** Spot tests. Indicated strains were transformed with an empty vector or the pREP41-GFP-HSET plasmid. Transformants were spotted on minimal plates in the presence of thiamine and incubated at indicated temperatures for 3 d. *cell conc*., cell concentration, *temp*., temperature. **(C)** Forced tethering of GFP-HSET to the SPB in wild-type or *pkl1*Δ cells. GFP-HSET (integrated at the *lysl* locus, see Materials and Methods) was recruited to the SPB using the same strategy as in Fig. 5C. Representative images of GFP-HSET during mitosis are shown. SPB-tethered GFP-HSET signals are marked with arrowheads. Scale bar, 10 μm. **(D)** Rescue of spindle anchoring defects in *pkl1*Δ cells by forced tethering of HSET to the SPB. >25 mitotic cells were observed in each strain. All p-values are derived from a two-tailed χ^2^ test (*, P < 0.05; ***, P < 0.001). **(E)** Growth inhibition by overproduced Pkl1, Klp2 or HSET. Wild-type cells containing a microtubule marker (mCherry-Atb2; red) were transformed with an empty vector (top row) or plasmids containing *pkl1* (pREP41-GFP-Pkl1, second row), *klp2* (pREP1-Klp2, third row) or *HSET* (pREP41-GFP-HSET, bottom row). Transformants were spotted onto minimal plates in the presence (left) or absence (right) of thiamine and incubated at 30°C for 3 d. *cell conc*., cell concentration. **(F)** Spindle morphology. Cells used in **(E)** were grown in minimal liquid media in the absence of thiamine for 22 h at 30°C. Cells contain a microtubule marker (mCherry-Atb2; red) and an SPB marker (Pcp1-CFP, blue) (Fong et al., 2010). Representative mitotic images are shown. Locations of mitotic SPBs (Pcp1-CFP) are marked with arrowheads. >80% cells (n>30) producing GFP-Pkl1, Klp2 or GFP-HSET displayed monopolar spindles with unseparated SPBs. Scale bar, 10 μm.

Pkl1 is localized to the mitotic SPB, while HSET was mainly localized along the spindle microtubule (Figs. 5A,6A), which might be the reason for inability of HSET to anchor the spindle microtubule as in the case of Klp2. We, therefore, tethered GFP-HSET to the SPB by implementing the same strategy as described earlier (GBP-Alp4). Tethering was successful (Fig. 6C) and intriguingly, SPB-localizing HSET indeed rescued the anchoring defects of *pkl1*Δ cells (Fig. 6D, n>25). Therefore, as in Klp2, it is the cellular localization rather than motor specificity that defines the functional differentiation between Pkl1 and HSET. We also tethered Cut7-GFP to the SPB using GBP-Alp4 in *pkl1*Δ cells (Fig. S2A) and examined spindle anchorage. In this case, unlike Kinesin-14s, anchoring defects were not suppressed (Fig. S2B). This result implies that despite that different Kinesin-14 members are operational in anchoring the spindle microtubule at the SPB, the type of kinesins is crucial as the plus-end directed motor is non-functional.

It is known that ectopic overproduction of Pkl1 blocks SPB separation through disproportionate inward force, resulting in the appearance of monopolar spindle phenotypes (Pidoux et al., 1996). We addressed whether other Kinesin-14s also have the same impact when overexpressed. We found that overexpression of *klp2* or *HSET* gene driven by the *nmt1* or *nmt41* promoter, respectively (Basi et al., 1993; Maundrell, 1990), inhibited cell growth and importantly, resulted in mitotic arrest with monopolar spindles (Fig. 6E,F, n>30), virtually identical to those imposed by Pkl1 overproduction or Cut7 inactivation (Hagan and Yanagida, 1990; Pidoux et al., 1996). Therefore, the three Kinesin-14s are capable of generating inward force to a similar degree that antagonizes outward force produced by Kineisn-5/Cut7.

## DISCUSSION

Fission yeast contains two Kinesin-14s, Pkl1 and Klp2, that generate inward force against outward force elicited by Kinesin-5/Cut7. Our study has uncovered interplay between Pkl1 and Klp2 that are localized to distinct mitotic structures. Pkl1 is localized to the mitotic SPB, thereby ensuring the anchorage of the spindle microtubule to the SPB, leading to the production of inward force. In contrast, Klp2 is localized along the spindle microtubule, where this motor generates pulling force by crosslinking antiparallel microtubules and sliding them inward. Notably, SPB-tethered Klp2 can replace the role of Pkl1 in spindle anchorage and force generation. Human Kinesin-14 HSET compensates for the loss of either Pkl1 or Klp2. We propose that it is the cellular localizations rather than individual motor specificities that determine the functional differentiation of each Kinesin-14 member.

### Force generation by two Kinesin-14 Pkl1 and Klp2

Although deletion of either *pkl1* or *klp2* is capable of rescuing various *cut7* ts mutants, only *pkl1*Δ can bypass the requirement of Cut7; *cut7*Δ*pkl1*Δ cells, but not *cut7*Δ*klp2*Δ cells, are viable. We envisage that provided the microtubule minus end is anchored to the SPB by the Pkl1 complex (MWP), cells lacking Cut7 and Klp2 cannot split duplicated SPBs due to the existence of Pkl1-mediated inward force. In contrast, in the absence of Pkl1, although Klp2-mediated inward force is exerted on the spindle microtubule, their minus ends are not tethered to the SPB, by which compromised inward force allows SPB separation. Alternatively, Klp2, which is localized to the overlapping spindle midzone, may not be able to generate robust inward force during the early mitotic stage (see below).

How do Pkl1 and Klp2 collaborate with each other to promote and maintain spindle bipolarity antagonistically with Cut7? We consider that the importance of Pkl1 and Klp2 may differ in each mitotic stage. In early mitotic stage, when spindle bipolarity starts to be established as the two SPB separate, Pkl1, which is localized to one SPB, engages with the spindle microtubule emanating from the opposite SPB. Under this condition, SPB-tethered Pkl1 walks towards the SPB, by which this minus-end motility produces pulling inward force that antagonizes Cut7-mideated outward force (Fig. 7A). To convert minus-end motility to inward force, the physical interaction between Pkl1 and the SPB (the γ-tubulin complex, γ-TuC) is essential (Olmsted et al., 2013; Rodriguez et al., 2008; Yukawa et al., 2015). During this stage, the involvement of Klp2 may be marginal, if at all. This is why *cut7*Δ*pkl1*Δ, but not *cut7*Δ*klp2*Δ, is viable. On the other hand, during the later mitotic stage, Klp2, which is localized to the spindle midzone, generates inward force in a manner opposing to Cut7 that generates outward force (Fig. 7B). The emergence of abnormally long anaphase B spindles in the absence of Klp2 is consistent with this notion (Troxell et al., 2001). Whether SPB-localizing Pkl1 produces inward force during this stage is not known; however, Pkl1 plays an important role in spindle anchorage, which resists Cut7-driven outward force as a barrier (Yukawa et al., 2015).

**Figure 7.**
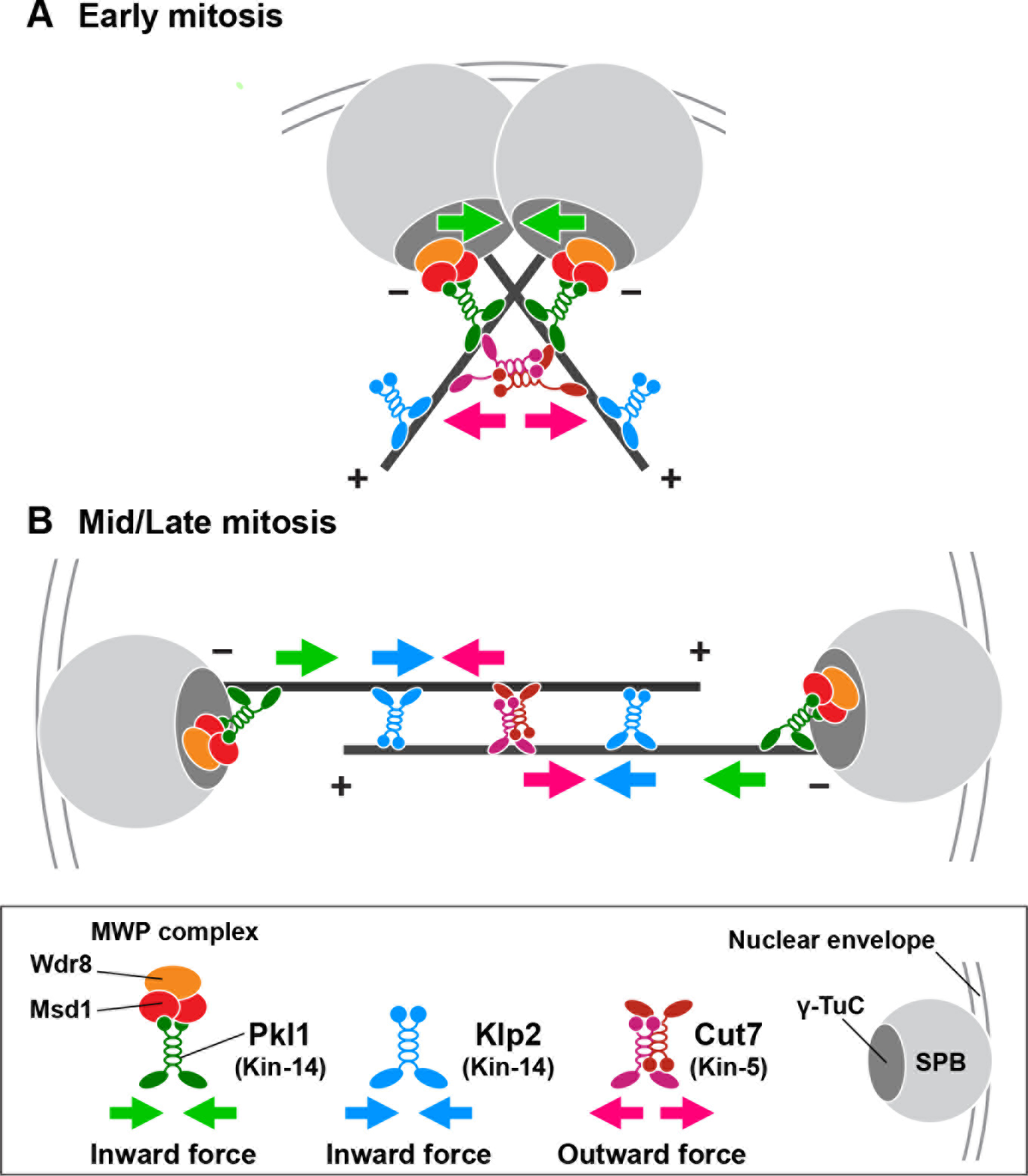
A model illustrating generation of collaborative inward forces by Pkl1 and Klp2. **(A)** During early mitosis, spindle microtubules nucleate from both SPBs and display a V-shaped morphology. Pkl1 is localized to the SPBs by forming a ternary complex with Msd1 and Wdr8 through binding to the the γ-tubulin complex (γ-TuC) (Olmsted et al., 2013; Rodriguez et al., 2008; Yukawa et al., 2015). SPB-tethered Pkl1 engages with the spindle microtubule that emanates from the opposite SPB. Minus end-directed motility of Pkl1 generates inward force (green arrows). This pulling force antagonizes outward force exerted by Cut7 (red arrows) that is localized to the spindle region in which parallel microtubules interconnect. It is not clear at the moment whether Klp2 generates force during this stage. Cut7 might also somehow inhibit Pkl1 as previously reported (Olmsted et al., 2014). **(B)** During the later stage of mitosis, antiparallel microtubules are formed in the spindle midzone, where Cut7 and Klp2 are localized. These two kinesins antagonistically generate outward force (red arrows) and inward force (blue arrows) respectively. Whether SPB-localizing Pkl1 generates inward force during this stage is unknown; however it is shown that Pkl1 plays a crucial role in anchoring the minus end of the spindle microtubules, thereby resisting Cut7-mediated outward force as a barrier (green arrows) (Yukawa et al., 2015). For simplicity, other microtubule-binding proteins including the microtubule crosslinker Ase1, the microtubule stabilizer Cls1/Peg1/CLASP, Kinesin-6 Klp9 and the microtubule polymerase complex Alp7/TACC and Alp14/T0G (Rincon et al., 2017; Yukawa et al., 2017) are not included in this figure.

### Kinesin-14s in yeasts, plants and humans

Our functional analysis indicates that both Pkl1 and Klp2 are deemed to be homologs of HSET, as this human motor protein, when introduced into fission yeast, can generate inward force in place of either of these two fungal Kinesin-14s. In human cells, HSET accumulates at the centrosome through interaction with centrosomal CEP215/CDK5RAP2 and/or the γ-tubulin complex (Cai et al., 2010; Chavali et al., 2016). This localization pattern mirrors that of Pkl1, as it is tethered to the SPB through a direct interaction between Pkl1 and γ-tubulin (Olmsted et al., 2013; Rodriguez et al., 2008) and/or through binding between Msd1-Wdr8 and Alp4/GCP2, a core component of the γ-tubulin complex (Toya et al., 2007; Yukawa et al., 2015). Like HSET, Pkl1 possesses minus end-directed motility (DeLuca et al., 2001; Furuta et al., 2008). In contrast, the mitotic localization of Klp2 is more complex, as it is localized along the lattice of spindle microtubules (Troxell et al., 2001) (and this study), to the microtubule plus end (Mana-Capelli et al., 2012; Scheffler et al., 2015) and the kinetochore (Gachet et al., 2008; Grishchuk and McIntosh, 2006; Troxell et al., 2001). Accordingly, the Klp2 motor appears to play multiple roles in addition to that in generation of inward force along the spindle microtubules. These notions indicate that Pkl1, rather than Klp2, is functionally more similar to HSET.

However, there are several features that are shared by Klp2 and HSET, but not by Pkl1, including microtubule-bundling activity (Braun et al., 2009; Cai et al., 2009; Carazo-Salas et al., 2005; Carazo-Salas and Nurse, 2006; Daga et al., 2006), a direct interaction with the microtubule plus end tracking proteins EB1/Mal3 (Braun et al., 2013; Mana-Capelli et al., 2012; Scheffler et al., 2015) and kinetochore localization (Xu et al., 2014). We ponder the idea that fission yeast has evolved to be equipped with two spatially distinct Kinesin-14s, both of which retain some functional features of human HSET. Yet, Pkl1 and Klp2 may have undergone further diversifications, thereby executing fission yeast-specific roles such as the formation of the MWP complex and the organization of interphase microtubules respectively (Carazo-Salas et al., 2005; Carazo-Salas and Nurse, 2006; Daga et al., 2006; Yukawa et al., 2015). It should be noted that the human genome contains three more genes coding for additional Kinesin-14s (KifC2, KifC3 and Kif25) that are classified as the Kinesin-14B subfamily (Hirokawa et al., 2009). Functional analysis of these motors has started to emerge (Decarreau et al., 2017). It would be of great interest to see how these additional members contribute to generation of inward force in collaboration with HSET.

Plants also contain multiple Kinesin-14 members and have been shown to play distinct roles (Yamada and Goshima, 2017; Yamada et al., 2017). However, it is not known which members of Kinesin-14s generate inward force against Kinesin-5 and hence, further characterization is awaited. In budding yeast, a single Kar3 motor forms a hetero-complex with either Cik1 or Vik1 (Manning and Snyder, 2000). Interestingly, while Vik1 is required for Kar3 localization to the SPB, Cik1 is needed for Kar3 to be localized to the microtubule lattice, thereby crosslinking microtubules. It, therefore, appears that budding yeast has evolved to create two different binding partners for Kar3, by which Kar3 displays two separate localizations and functions; Kar3-Vik1 exhibits functional similarity to Pkl1, while Kar3-Cik1 is more analogous to Klp2. Collectively, we conclude that outward force generated by Kinesin-5 is balanced by dual inward forces exerted by different Kinesin-14s that are subject to distinct spatial regulation.

## MATERIALS AND METHODS

### Strains, media and genetic methods

Fission yeast strains used in this study are listed in Table S1. Media, growth conditions and manipulations were carried out as previously described (Bähler et al., 1998; Moreno et al., 1991; Sato et al., 2005). For most of the experiments, rich YE5S liquid media and agar plates were used. Wild-type strain (513; Table S1), temperature-sensitive *cut7* (*cut7-21*, *-22*, *-23*, *-24*, *-446*), *pkl1* deletion strains were provided by P. Nurse (The Francis Crick Institute, London, England, UK), I. Hagan (Cancer Research UK Manchester Institute, University of Manchester, Manchester, England, UK) and R. McIntosh (University of Colorado, Boulder, CO) respectively. For overexpression experiments using *nmt* series plasmids, cells were first grown in Pombe Glutamate Medium (PMG, the medium in which the ammonium of EMM2 is replaced with 20 mM glutamic acid) with required amino acid supplements in the presence of 15 μM thiamine overnight. Thiamine was then washed out by filtration pump and cells cultured in the same PMG media in the absence of thiamine for further 12~24 h as necessary. Spot tests were performed by spotting 5-10 μl of cells at a concentration of 2 × 10^7^ cells/ml after 10-fold serial dilutions onto rich YE5S plates or PMG plates with added supplements with or without 15 μM thiamine. Some of the YE5S plates also contained Phloxine B, a vital dye that accumulates in dead or dying cells and stains the colonies dark pink due to a reduced ability to pump out the dye. Plates were incubated at various temperatures from 27°C to 37°C as necessary.

### Preparation and manipulation of nucleic acids

Enzymes were used as recommended by the suppliers (New England Biolabs Inc. Ipswich, MA, U. S. A. and Takara Bio Inc., Shiga, Japan).

### Strain construction, gene disruption and the N-terminal and C-terminal epitope tagging

A PCR-based gene-targeting method (Bähler et al., 1998; Sato et al., 2005) was used for complete gene disruption and epitope tagging (e.g., GFP, CFP and mCherry) in the C terminus, by which all the tagged proteins were produced under the endogenous promoter. A strain containing GFP-Pkl1 or GFP-Klp2 was constructed as follows. DNA fragments containing a G418-resistance gene (*kan*), the *alp4* promoter and *GFP* (Masuda et al., 2013) were PCR-amplified and inserted in-frame to the 5’-franking region of the *pkl1*^+^ or *klp2*^+^ *ORF* by the fusion PCR method. GBP-mCherry-Alp4 was constructed as follows. DNA fragments containing a G418-resistance gene (*kan*), the *alp4* promoter, *GBP* and *mCherry* were PCR-amplified and inserted into the 5’-flanking region of the *alp4 ORF* in-frame by the fusion PCR method. To construct a strain carrying GFP-HSET, the *HSET* cDNA (DNAFORM Inc., Clone ID: 40117894, Yokohama, Japan) was amplified with a pair of primers carrying *Sal*I and *Bam*HI sites and the PCR product was ligated in-frame to the 3′ end of *GFP* at the *Xho*I/*Bgl*HI sites of pCSU71 (Chikashige et al., 2004). The resulting plasmid was integrated into the *lysl-l* locus, which produces Lys^+^ colonies.

### Fluorescence microscopy and time-lapse live cell imaging

Fluorescence microscopy images were obtained using a DeltaVision microscope system (DeltaVision Elite; GE Healthcare, Chicago, IL, U. S. A.) comprising a wide-field inverted epifluorescence microscope (IX71; Olympus, Tokyo, Japan) and a Plan Apochromat 60×, NA 1.42, oil immersion objective (PLAPON 60×0; Olympus Tokyo, Japan). DeltaVision image acquisition software (softWoRx 6.5.2; GE Healthcare, Chicago, IL) equipped with a charge-coupled device camera (CoolSNAP HQ2; Photometrics, Tucson, AZ) was used. Live cells were imaged in a glass-bottomed culture dish (MatTek Corporation, Ashland, MA) coated with soybean lectin and incubated at 27°C for most of the strains or at 36°C for the ts mutants. The latter were cultured in rich YE5S media until mid-log phase at 27°C and subsequently shifted to the restrictive temperature of 36°C before observation. To keep cultures at the proper temperature, a temperature-controlled chamber (Air Therm SMT; World Precision Instruments Inc., Sarasota, FL) was used. The sections of images acquired at each time point were compressed into a 2D projection using the DeltaVision maximum intensity algorithm. Deconvolution was applied before the 2D projection. Images were taken as 14-16 sections along the z axis at 0.2 μm intervals; they were then deconvolved and merged into a single projection. Captured images were processed with Photoshop CS6 (version 13.0; Adobe, San Jose, CA).

### Statistical data analysis

We used the two-tailed unpaired Student’s *t*-test to evaluate the significance of differences in different strains. All the experiments were performed at least twice. Experiment sample numbers used for statistical testing were given in the corresponding figures and/or legends. We used this key for asterisk placeholders to indicate p-values in the figures: e.g., ****, P < 0.0001.

## Acknowledgements

We are grateful to Yuji Chikashige, Iain Hagan, Yasushi Hiraoka, Heinrich Leonhardt, J. Richard McIntosh, Paul Nurse and Gislene Pereira for providing us with strains used in this study. We thank Ken’ya Furuta for thoughtful discussion and Corinne Pinder and Richard O’Connor for critical reading of the manuscript and useful suggestions. We are grateful to Ayaka Inada for her technical assistance.

## Author contributions

Conceptualization: Y.M., T.T.; Methodology: M.Y., Y.Y., T.Y; Formal analysis: M.Y., Y.Y., T.Y.; Resources: Y.M., T.T.; Writing - original draft: Y.M.; Writing - review & editing: Y.M., T.T.; Supervision: Y.M., T.T.; Project administration: Y.M., T.T.; Funding acquisition: Y.M., T.T.

## Supplementary information

Supplementary information is available online.

